# Effects of early life stress and subsequent re-exposure to stress on neuronal activity in the lateral habenula

**DOI:** 10.1101/2022.08.11.503558

**Authors:** Jack F. Webster, Sanne Beerens, Christian Wozny

## Abstract

Early life stress can result in depression in humans and depressive-like behaviour in rodents. In various animal models of depression, the lateral habenula (LHb) has been shown to become hyperactive immediately after early life stress. However, whether these pathological changes persist into adulthood is less well understood. Hence, we utilised the maternal separation (MS) model of depression to study how early life stress alters LHb physiology and depressive behaviour in adult mice. We find that only a weak depressive phenotype persists into adulthood which surprisingly is underpinned by LHb hypoactivity in acute slices, accompanied by alterations in both excitatory and inhibitory signalling. However, while we find the LHb to be less active at rest, we report that the neurons reside in a sensitised state where they are more responsive to re-exposure to stress in adulthood in the form of acute restraint, thus priming them to respond to aversive events with an increase in neuronal activity mediated by changes in glutamatergic transmission. These findings thus suggest that in addition to LHb hyperactivity, hypoactivity likely also promotes an adverse phenotype. Re-exposure to stress results in the reappearance of LHb hyperactivity offering a possible mechanism to explain how depression relapses occur following previous depressive episodes.

## Introduction

The lateral habenula (LHb) is an evolutionarily conserved brain structure located within the epithalamus which encodes aversive events (Matsumoto and Hikosaka, 2007; Lecca *et al*., 2017), and depressive behaviour (Li *et al*., 2011; Yang *et al*., 2018; Hu *et al*., 2020; Zheng *et al*., 2022). Specifically, the LHb becomes hyperactive in depression (Li *et al*., 2011; Lecca *et al*., 2016; Tchenio *et al*., 2017; Cui *et al*., 2018; Yang *et al*., 2018), thus enhancing output to the midbrain reward circuitry, for which the LHb acts as an inhibitory modulator (Wang and Aghajanian, 1977; Ji and Shepard, 2007; Jhou *et al*., 2009).

Indeed, many studies have employed the use of a variety of different animal models of depressive behaviour including chronic mild stress (Cerniauskas *et al*., 2019), chronic restraint stress (Yang *et al*., 2018; Zheng *et al*., 2022), social defeat (Golden *et al*., 2016; Knowland *et al*., 2017), learned helplessness (Li *et al*., 2011) and various models of early life stress (Tchenio *et al*., 2017; Authement *et al*., 2018; Simmons *et al*., 2020; Langlois *et al*., 2022), and have independently reached this conclusion that the LHb becomes hyperactive in depression. However, the majority of these studies have carried out experimentation shortly after exposure to the relevant stressor, and as such there is comparatively little evidence as to the long-term persistence of depressive phenotype, and the corresponding synaptic and physiological alterations within the LHb (Langlois *et al*., 2022).

Hence, in this study we aimed to assess how early life stress in the form of maternal separation influences depressive behaviour, and alters lateral habenular physiology and synaptic connectivity in adult mice. Furthermore, we then sought to ask how early life stress influences subsequent re-exposure to stress in adulthood.

## Materials and methods

### Animals and maternal separation procedure

All procedures were approved by the Ethics committee of the University of Strathclyde, Glasgow, and carried out in accordance with the relevant UK legislation (the Animals (Scientific Procedures) Act, 1986). Male and female mice from each strain were used in this work, and unless otherwise stated, data were pooled between genders. Mouse strains used in this study were C57BL/6J, SOM-IRES-Cre heterozygous mutants (Jax ID 018973; Taniguchi *et al*., 2011), and their wild-type littermates. All animals were kept on a 12:12 light/dark cycle under standard group housing conditions with unlimited access to water and normal mouse chow, unless otherwise stated.

The maternal separation (MS) procedure was adapted from a previously published protocol (Tchenio *et al*., 2017). Pregnant females were housed individually, and date of littering was counted as postnatal day 0 (P0). Litters of 5-10 pups were used for these procedures. At P6, the litter was divided into even (±1 pups) groups of MS and control (CTRL) pups. MS pups were then separated from the mother into individually isolated compartments in a heated cage in a separate room for 6 hours per day from P6-16, and then weaned early at P17. Separation started each day between 08:00 and 10:00. CTRL pups remained with the mother and were weaned at P21. At the early weaning age of the MS pups (P17), they are often unable to consume normal mouse chow, and as such we provided them with human baby food for the final 2 days of MS to allow them to habituate to this. This was then provided for several days post weaning for both MS and CTRL mice to allow them to rapidly develop to the point where they could be sustained on normal mouse chow. Following weaning, both groups were then group housed (2-5 mice per cage) and allowed to develop to adulthood when further testing commenced.

### Behavioural testing and acute restraint procedure

Mice underwent behavioural testing at approximately 8-10 weeks, and were single housed for these experiments. 3 behavioural tests were implemented in this study. These were sucrose preference, the open field test, and the splash test. Mice were then returned to group-housing conditions upon completion of behavioural testing.

#### Sucrose preference

Mice were first single housed, and then were given a choice between 2 bottles of water to habituate them to having 2 spouts in the cage. The following day (Day 1), both bottles were refilled and replaced. The left bottle again contained water, and the right bottle was filled with 1 % sucrose solution. On the third day (Day 2), bottles were refilled and replaced, this time with the 1 % sucrose solution bottle on the left, and water on the right. On the final day (Day 3), the positions of the bottles were pseudorandomised such that they alternated between cages. For all days, bottles were weighed before being added to the cage and again 24 hours later, and the consumption of fluid from each was used to calculate sucrose preference.

#### Open field test

Open field testing was carried out the day following the completion of sucrose preference testing. Mice were put in a white square arena (40 × 40 × 35 cm) within a room with standard lighting conditions, and allowed to freely explore for 5 minutes. Activity was recorded using an overhead camera, and videos were analysed using the analysis programme ToxTrac (Rodriguez *et al*., 2018), with the outermost 5 cm of the arena being classed as the borders.

#### Splash test

Splash test was carried out on the same day as open field testing, at least 2 hours after completion of this test. This was carried out in the home cage. 30 minutes prior to testing, all environmental enrichment (nesting material and plastic house) was removed from the home cage. Mice were then sprayed once on the dorsal coat with a solution of 10 % sucrose. To avoid the scent of sucrose distracting the animals during testing, mice were sprayed outside of the home cage. The mouse was then immediately returned to the home cage, the lid replaced, and activity recorded for 5 minutes using an overhead camera. Analysis was performed using the analysis programme BORIS (Friard and Gamba, 2016).

#### Acute restraint

Mice were restrained within modified handling tubes for a period of 1 hour. Upon completion of the acute restraint procedure, mice were either immediately sacrificed by cervical dislocation for preparation of acute brain slices, or returned to the home cage for 1 hour before transcardial paraformaldehyde (PFA) perfusion to assess c-Fos expression. Mice that were to be sacrificed for brain slice preparation immediately after restraint remained in group-housing conditions. Mice that were to undergo PFA perfusion were single-housed for at least 2 days prior to restraint.

#### Behavioural Z scoring

Behavioural Z scores were calculated as previously described (Guilloux *et al*., 2011). Briefly, a normalised Z score was calculated for each of the following 7 behavioural readouts: sucrose preference on days 1, 2 and 3; percentage time spent in borders in the open field test and; percentage time spent grooming, latency to first grooming bout and time rearing in the splash test. Behavioural Z scores were then calculated as the average of these 7 Z scores for each individual animal.

### Stereotaxic viral injections

SOM-IRES-Cre heterozygous mice (approximately 8-9 weeks old) were deeply anaesthetized via inhaled isoflurane (5% for induction; 1–2% for maintenance), transferred to a stereotaxic frame (Narishige, Tokyo, Japan) and were subcutaneously injected with the analgesics carprofen (5 mg/kg) in the nape and lidocaine (4 mg/kg) under the scalp. Intracranial injections were made using a glass micropipette pulled using a PC-100 vertical puller (Narishige, Tokyo, Japan). Under aseptic conditions, the skull was exposed and a small burr hole was drilled bilaterally above the basal forebrain (BF). Stereotaxic coordinates (from Bregma) were as follows: AP 0.45; ±1.3; depth 5.8. The injection capillary was then advanced and viral vector solutions were injected at a rate of 100 nL/min using a pressure microinjector (Narishige, Tokyo, Japan). Viral vector solutions used in this study were AAV9-EF1a-DIO-hChR2(H134R)-EYFP, titre 1.8×10^13^ vg/mL, 200 nL; AAV9-pCAG-FLEX-EGFP-WPRE, titre 2.5×10^13^ vg/mL, 100 nL injected (both from Addgene, Massachusetts, US). Following injection, the needle was left for at least 10 minutes to allow the virus to diffuse before being slowly withdrawn. Animals were allowed to recover from anaesthesia on a heat pad. Following completion of surgery, animals were given at least two weeks to allow expression of the virus before acute slice preparation for electrophysiology. Assessment of viral spread was carried out using either a fluorescent camera (Olympus XM10; Olympus, Southend-on-Sea, UK) with a 4X objective, or a Leica SP8 confocal microscope.

### Acute brain slice preparation

Mice were humanely euthanized by cervical dislocation and immediately decapitated, and brains were rapidly removed and transferred to ice-cold oxygenated (95% O2; 5% CO2) sucrose-based artificial cerebrospinal fluid (ACSF) solution containing (in mM): sucrose 50, NaCl 87, NaHCO3 25, KCl 3, NaH2PO4 1.25, CaCl2 0.5, MgCl2 3, sodium pyruvate 3 and glucose 10. Brains sections containing the lateral habenula were then cut in the coronal plane at 250 µm where *in vitro* optogenetic experiments were to be performed, or 300 µm for all other slice experiments, on a Leica VT1200S vibratome (Leica Biosystems, Newcastle-upon-Tyne, UK). Following sectioning, slices were incubated in oxygenated sucrose-based ACSF at 35 °C for 30 minutes, and then incubated for a further 30 minutes at room temperature in ACSF containing (in mM) NaCl 115, NaHCO3 25, KCl 3, NaH2PO4 1.25, CaCl2 2, MgCl2 1, sodium pyruvate 3 and glucose 10. Following the incubation period, slices were stored at room temperature in oxygenated ACSF until required for electrophysiological recordings.

### *In vitro* electrophysiological recordings

Individual slices were transferred to a recording chamber and continually perfused with oxygenated ACSF at a flow rate of 2–3 mL/min and visualized with a Luigs and Neumann LN-Scope System (Luigs and Neumann, Ratingen, Germany). Neurons suitable for whole-cell recordings were identified under a 60X objective. For transgenic animals which expressed a fluorescent reporter protein (eGFP or eYFP), fluorescent reporters were excited using a LED coupled through the 60X objective (pE-300ultra, Cool LED, Andover, UK), and reporter-expressing somata or terminal fields were visualized with an Olympus XM10 fluorescent camera (Olympus, Southend-on-Sea, UK)). Recordings were made with a Multiclamp 700B Amplifier (Molecular Devices, California, USA). For current clamp recordings, and for recording optogenetically-driven postsynaptic currents, glass micropipettes were filled with a solution containing (in mM) potassium gluconate 125, Hepes 10, KCl 6, EGTA 0.2, MgCl2 2, Na-ATP 2, Na-GTP 0.5, sodium phosphocreatine 5, and with 0.2% biocytin, and pH was adjusted to 7.2 with KOH. For spontaneous postsynaptic current measurement experiments, a potassium chloride-based intracellular solution was used consisting of (in mM) potassium chloride 145, EGTA 0.1, Hepes 10, NaATP 2 and MgCl2 2, pH adjusted to 7.2. To pharmacologically isolate inhibitory currents, these experiments were performed in the presence of AMPA and NMDA receptor blockade (10 µM NBQX and 50 µM D-AP5 respectively; Tocris, Bristol, UK). To isolate excitatory currents, experiments were performed in the presence of GABAA and GABAB blockade (5 µM SR-95531 and 10 µM CGP-52432 respectively; Tocris, Bristol, UK).

For current clamp recordings, a gigaseal was first achieved in voltage-clamp configuration when the pipette was in the immediate vicinity of the neuron. The neuron was then held at a potential of -60 mV, and whole-cell configuration was achieved by rupturing the membrane with a series of negative pressure pulses. Once in whole-cell patch mode, the intrinsic properties of LHb neurons were assessed by switching to current-clamp configuration (*I = 0)* and recording spontaneous activity for a period of 2-3 minutes. Neurons which fired action potentials with a frequency of at least 0.5 Hz were classed as spontaneously active. Following spontaneous activity recording, current-spike input-output relationships were tested by injecting sufficient holding current to hold the neuron at a potential of -55 mV, and a series of depolarising current steps were injected (0-100 pA; 10 pA steps). Holding current was then removed, and a second series of current steps were injected (−50-100 pA; 10 pA steps) to assess both rebound bursting properties, and spiking properties at rest.

For optogenetically-driven postsynaptic inhibitory current recordings, whole-cell configuration was achieved as above, and recordings were performed in voltage-clamp configuration at a holding potential of -50 mV. During recording, a single blue LED pulse (1 ms) of increasing intensity (1-50%; 3-4 trials per intensity) was applied to induce a postsynaptic current, with the amplitude being measured as an average of 3 trials. Upon completion of these experiments, the same neurons were then held in current clamp configuration to assess both physiological properties, and the capacity for optogenetically-driven inhibitory synaptic transmission to induce rebound bursting.

For spontaneous current measurements, whole-cell configuration was achieved at -70 mV, and spontaneous synaptic activity was recorded for a period of 2 minutes. For these experiments, recordings were completed no more than 4 minutes after break-in, as we observed rapid reduction in synaptic event frequency using the aforementioned high chloride intracellular solution.

Series resistance was monitored throughout. All neuronal voltage and current signals were low pass-filtered between 2 and 10 kHz and acquired between 10 and 25 kHz using an ITC-18 digitizer interface (HEKA, Pfalz, Germany). The data acquisition software used was Axograph X.

### Transcardial perfusion and sectioning

Mice were terminally anaesthetized by intraperitoneal injection with an overdose cocktail of 50% lidocaine and 50% pentobarbital. Once anaesthetized sufficiently to be non-responsive to tail and toe pinch stimuli, mice were perfused through the left ventricle with 0.1 M PBS followed by perfusion with 4% PFA dissolved in PBS. Brains were then removed and fixed overnight in 4% PFA in PBS, after which they were cryoprotected in a solution containing 30% sucrose in PBS. Brains were left in this solution until they dropped to the bottom of the tube, at which point the 30% sucrose solution in PBS was replaced with fresh 30% sucrose solution. Once the brain dropped for a second time, it was considered ready for sectioning. This was performed by embedding in OCT compound (VWR, Leicestershire, UK) and freezing with a dry ice bath. Once frozen, brains were sectioned on a Leica SM2010 R microtome (Leica Biosystems, Newcastle-upon-Tyne, UK) at 50 - 60 µm.

### Immunohistochemistry and confocal imaging

Following sectioning, slices were washed 3 times in 0.1 M PBS, and then incubated for 30 minutes in a blocking solution consisting of 5% normal goat serum (NGS) and 0.3% Triton X-100. Blocking solutions was then removed, and slices were incubated on a shaker at room temperature overnight in a primary antibody mixture containing 0.3% Triton in PBS and rabbit anti-c-Fos (1/10000; ab190289; Abcam, Cambridge, UK). Upon completion of the primary incubation step, slices were washed 2 × 5 minutes in 0.1 M PBS and incubated for 3 hours in a solution containing donkey anti-rabbit conjugated to Alexa Fluor 647 (1/500 dilution; Invitrogen, UK). After secondary antibody incubation, slices were washed for 3 times in 0.1 M PBS and mounted on glass slides using Vectashield medium containing DAPI (Vector Labs, Peterborough, UK).

Slices were imaged on a Leica SP8 confocal microscope using a 20X objective at 3 different points from Bregma along the rostrocaudal axis, spaced approximately 300 µm apart. For these experiments, imaging was performed with a 633 nm laser at 1 % max intensity, 2 µm z-steps.

### Statistical analysis

Statistical analysis was carried out in GraphPad Prism 9.3.1. For pairwise comparisons, an unpaired T test was used where at least one data set was found to be normally distributed (tested with a Shapiro-Wilk normality test). Where both sets of data failed a normality test, a Mann-Whitney test was used.

## Results

### Early life stress induces mild depressive symptoms in adult mice, and alters LHb physiology

We first sought to validate that MS for 6 hours per day with early weaning (methods) can induce depressive behaviour in adult mice. Hence, following MS, mice were allowed to develop to adulthood (≈ 8-10 weeks of age), when they underwent behavioural testing. Mice were subjected to a series of 3 behavioural paradigms to test for anhedonia, anxiety-like behaviour and motivation which were sucrose preference, the open field test and the splash test respectively (Fig. 1A; N = 37 CTRL; N = 39 MS mice). Sucrose preference testing was carried out over 3 consecutive days, and indeed we saw a reduction in sucrose preference in MS mice over these 3 days (Fig. 1B; *p* = 0.038; 2-way ANOVA column factor). However, a particularly striking effect was that this deficit was most prominent on day 2 of testing (*p* = 0.028; Sidak’s test). This was an interesting observation, as on day 2 of our paradigm, we switched the position of the sucrose and water bottles (Methods), hence suggesting that our model also induces a possible deficit in reversal learning (Baker and Mizumori, 2017). We did not observe any change in anxiety-like behaviour (Fig. 1Ci; *p* = 0.670; unpaired T-test), locomotor activity (Fig. 1Cii; *p =* 0.593; unpaired T-test) or motivation (Fig. 1Di; *p* = 0.286; unpaired T-test). However, MS mice exhibited an increased latency to first grooming bout in the splash test (Fig. 1Dii; *p* = 0.050; Mann-Whitney test), and interestingly spent more time rearing in the splash test (Fig. 1Diii; *p* = 0.020; unpaired T-test), which may be indicative of social contact seeking (Fukumitsu *et al*., 2022). To account for variability between behavioural tests within individual mice, we also calculated an integrated Z score (Methods and Guilloux *et al*., 2011), which gives an arbitrary score of emotionality for each mouse, by normalising and integrating readouts from each behavioural test. This indeed revealed that MS mice had overall greater emotionality than CTRL mice (Fig. 1E; *p* = 0.019; unpaired T-test). However, overall these results led us to conclude that the observed phenotype was relatively mild in the adult mice.

**Figure 1:**
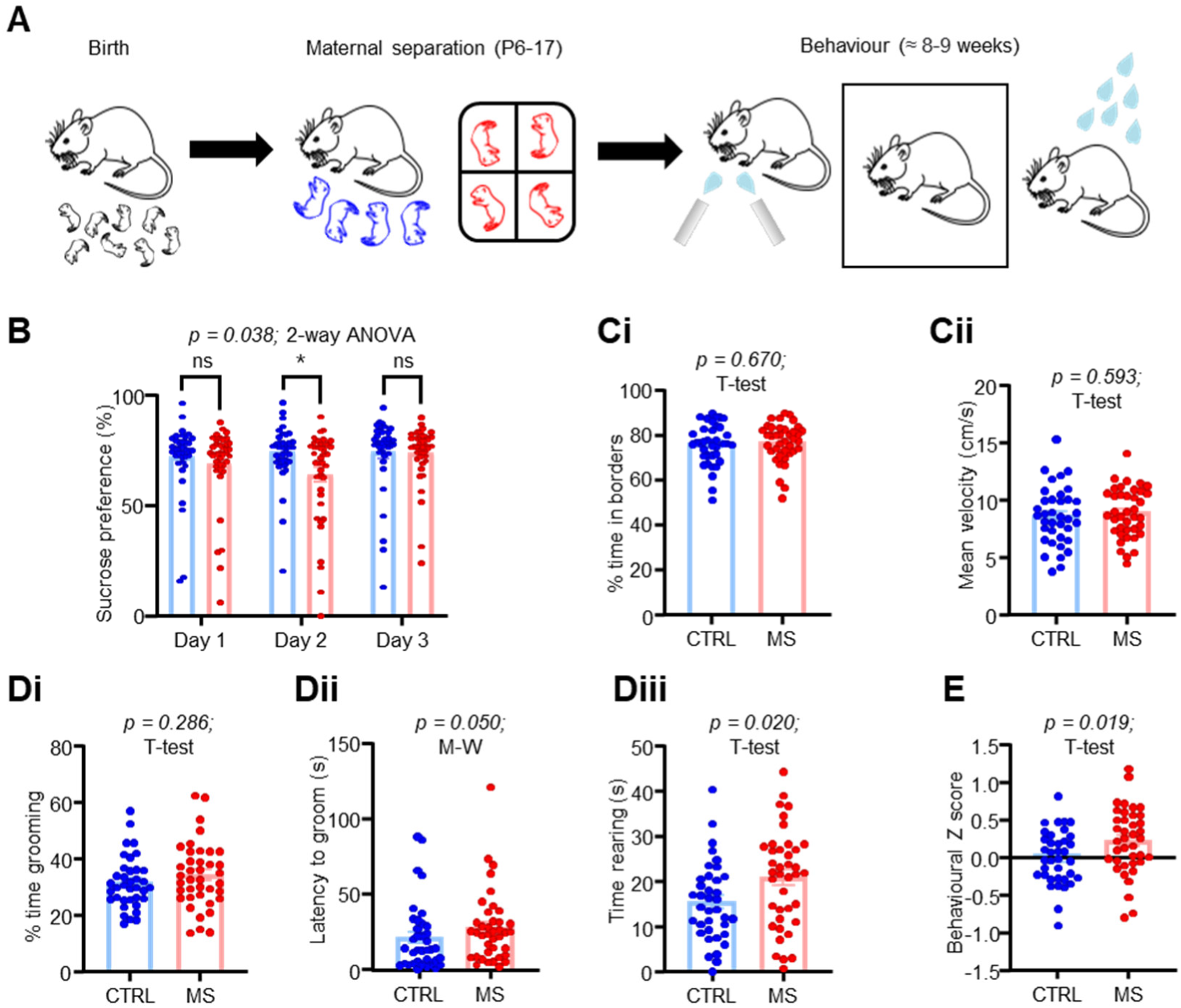
MS induces a mild depressive-like phenotype in adult mice. (A) Schematic illustrating experimental timeline. (B) Behavioural data from sucrose preference test across 3 days of testing. (C) Behavioural data from open field test. (Ci) Comparison of % of test time in borders and of (Cii) mean locomotor activity. (D) Behavioural data from splash test. (Di) Comparison plot of % of test time spent grooming; (Dii) of latency to first grooming bout and (Dii) of total time spent rearing on hind legs. (E) Averaged z scores of behavioural traits for each mouse.

We next sought to assess how MS altered LHb neuronal physiology. Mice were sacrificed shortly after behavioural testing, and whole-cell recordings were carried out in acute brain slices (Fig. 2A; n/N = 36/6 CTRL; n/N = 39/6 MS neurons / mice). MS induced no changes in passive physiological properties (Fig. S1A; input resistance *p* = 0.984; unpaired T-test; and resting membrane potential *p* = 0.235; Mann-Whitney test), and induced only a weak trend towards an increase in intrinsic excitability (Fig. S1B; *p* = 0.083; 2-way ANOVA column factor). However, the most striking difference we observed was a reduction in spontaneous neuronal activity (Fig. 2Bi and Bii; *p* = 0.0009; Chi-square test), which appeared to be specific for tonically active neurons. This was an interesting observation as bursting activity of LHb neurons is believed to be the primary driver of depressive behaviour (Yang *et al*., 2018; Zheng *et al*., 2022).

**Figure 2:**
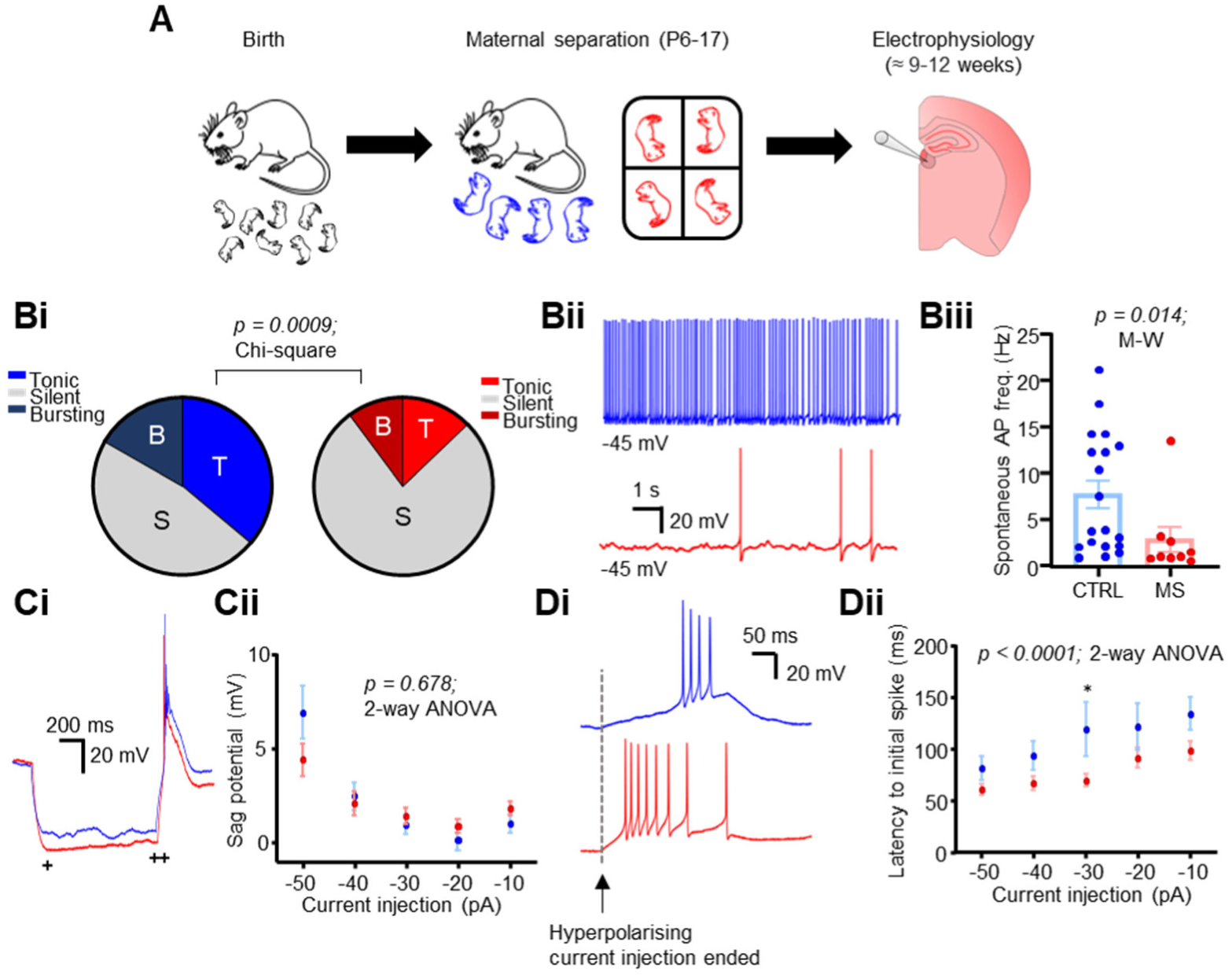
MS alters LHb neuronal physiology in adult mice. (A) Schematic illustrating experimental timeline. (B) Spontaneous activity comparison for both conditions. (Bi) Pie charts depicting fractions of recorded neurons which were classed as spontaneously active (spontaneous AP discharge > 0.5 Hz). (Bii) Example spontaneous activity recordings from both conditions. (Biii) Comparison plot of mean spontaneous activity frequency, for neurons which were spontaneously active (>0.5 Hz frequency). (Ci) Example traces and (Cii) plot of input current against sag potential in both conditions. Sag potential was calculated as the difference of the peak (+) and the steady state (++) of the membrane hyperpolarisation induced in response to hyperpolarising current steps. (Di) Example traces of rebound bursts induced in both conditions following hyperpolarising current injection. (Dii) Plot of input current against latency to initial spike of the rebound burst induced upon current step end.

Moreover, of those neurons which were spontaneously active, we observed that they were active at a lower frequency following MS (Fig. 2Bii; *p* = 0.014; Mann-Whitney test). We also observed no difference in sag potential between conditions (Fig. 2Ci and Cii; *p* = 0.678; 2-way ANOVA column factor). However, although neurons from MS mice did not display more bursting activity at rest, we found that they did fire rebound bursts with a reduced latency following hyperpolarising current injection (Fig. 2Di and Dii; *p* < 0.0001; 2-way ANOVA column factor), leading us to speculate that the neurons may be in a state where they are more primed to fire in response to synaptic input. Hence, we also recorded spontaneous excitatory postsynaptic currents (sEPSC’s) in a separate cohort of mice (Fig. S2; n/N = 44/4 CTRL; n/N = 46/4 MS neurons / mice). We did not observe any differences in frequency (Fig. S2Ai; *p* = 0.342; Mann-Whitney test) but did observe a slight but significant reduction in current amplitude following MS (Fig. S2Ai; *p* = 0.012; Mann-Whitney test). However, this dataset was likely confounded by the fact that the CTRL mice for this particular cohort appeared to display a phenotype more similar to the MS mice than other CTRL mice, and as such we attempted to account for this by plotting the behavioural Z score of each mouse against the mean sEPSC frequency for all cells recorded from each individual mouse (Fig. S2B). Indeed, when we did this we observed a negative correlation between Z score and mean sEPSC frequency (Fig. S2B; *p* = 0.016; simple linear regression), indicating that mice which existed in a more depressed state apparently exhibited reduced excitatory drive onto LHb neurons. Altogether, these results point to a scenario whereby MS reduces spontaneous firing of LHb neurons in brain slices, possibly by reducing presynaptic excitatory drive.

### MS weakens inhibitory synaptic transmission onto LHb neurons

Multiple other works have shown that a reduction in inhibitory signalling is associated with a depressive phenotype (Shabel *et al*., 2014; Lecca *et al*., 2016; Tchenio *et al*., 2017), and as such we sought to test how MS influenced inhibitory signalling in adult mice. We first recorded spontaneous IPSCs (sIPSCs) throughout the LHb in acute slices (Fig. 3; n/N = 46/6 CTRL; n/N = 49/6 MS neurons / mice). We observed no overall difference in either frequency (Fig. 3A; *p* = 0.296; Mann-Whitney test) or amplitude (Fig. 3A; *p* = 0.245; Mann-Whitney test). However, we did observe a striking gradient across the mediolateral axis of the LHb in CTRL mice (Fig. 3Bi and Bii; *p* = 0.005; simple linear regression) with sIPSC frequency being greatest in the medial LHb, which was not present in MS mice (Fig. 3Bi and Bii; *p* = 0.325; simple linear regression). This thus lead us to suspect that there may be subregional differences in spontaneous inhibitory synaptic input to LHb neurons. Indeed, when we broke our analysis down into the medial (< 0.45 mm from midline) and lateral (> 0.45 mm from midline) LHb sub-regions, we observed a reduction in sIPSC frequency specifically in the medial LHb (Fig. 3Bii; *p* = 0.048; Mann-Whitney test). Furthermore, we found that there was a negative correlation of the behavioural Z score with IPSC frequency (Fig. 3C; *p* = 0.013; simple linear regression), thus suggesting that more depressed animals exhibited lower levels of inhibitory synaptic drive. Overall, these results suggest that MS results in a reduction in spontaneous inhibitory input, specifically in the medial LHb.

**Figure 3:**
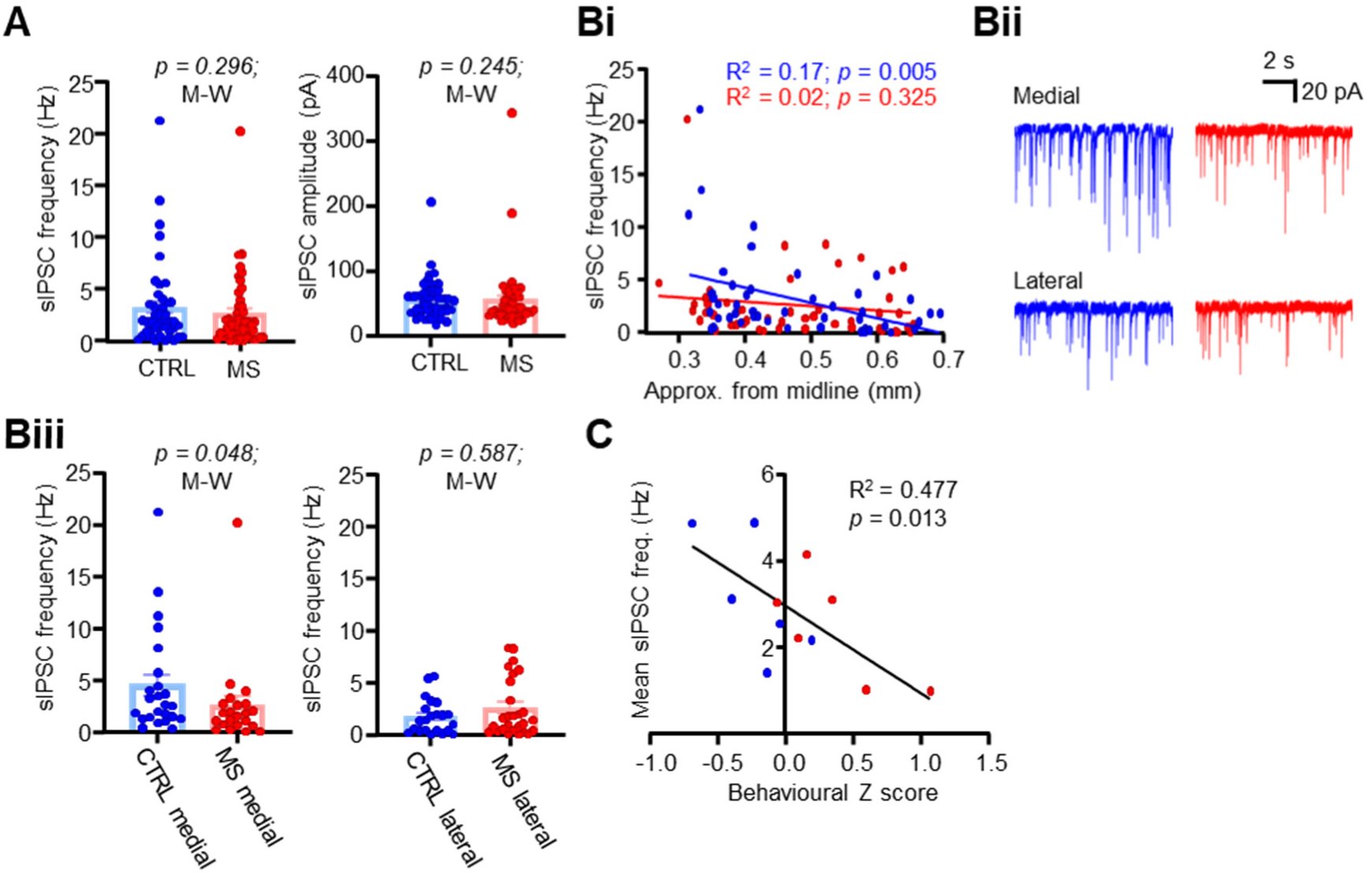
MS induces loss of spontaneous inhibitory input in the medial LHb. (A) Comparison plots of sIPSC frequency and amplitude between conditions. (Bi) XY plot with simple linear regression of sIPSC frequency against approximate distance from midline for both conditions. (Bii) Example sIPSc recordings from neurons recorded in both the medial and lateral LHb for both conditions. (Biii) Comparison plots of sIPSC frequency in both the medial and lateral LHb. (C) XY plot of behavioural z score against mean sIPSC frequency calculated for each individual mouse recorded from. Mean sIPSC scores are calculated as the mean sIPSC frequency of all cells recorded from each mouse.

In our previous work we reported on a strong inhibitory projection to the LHb which appeared to arise from somatostatin-positive (SOM) neurons in the ventral pallidum of the basal forebrain (Webster *et al*., 2020). Other work has shown that excitatory pallidal projections to the LHb can promote depressive behaviour (Knowland *et al*., 2017), and that inhibitory LHb-projecting pallidal projections promote reward (Stephenson-Jones *et al*., 2020). As such, we hypothesised that inhibitory drive onto the LHb from SOM-positive pallidal neurons may be lost following MS. We tested this by injecting a Cre-dependent virus encoding channelrhodopsin (ChR2) into the basal forebrain of SOM-Cre mice and recorded light-induced postsynaptic events in LHb neurons (Fig. 4A; n/N = 18/4 CTRL; n/N = 20/ 4 MS neurons / mice). Optogenetic stimulation induced inhibitory postsynaptic events in similar fractions of neurons in both CTRL and MS mice (Fig. 4B; *p* = 0.59; Chi square test). We first recorded inhibitory currents induced following a single 1 ms pulse at various intensities, and tested for differences between CTRL and MS mice by fitting a one-phase association exponential curve to each dataset (Libovner *et al*., 2020). CTRL and MS groups were found to have very different curve fits (Fig. 4Bi; *p* < 0.0001; non-linear curve fit), which specifically was found to be a reduction in the curve plateau for MS mice (Fig. 4Bi and Bii; *p* = 0.03; non-linear curve fit), thus suggesting a reduction in postsynaptic current amplitude without a corresponding change in kinetics. We also tested presynaptic release probability in current clamp configuration and observed no differences (Fig. 4C; *p* = 0.457, 2-way ANOVA). However, an interesting observation was that in a sub-fraction of responsive neurons, the inhibitory drive was found to be strong enough to induce rebound firing (Fig. S3A and B), which interestingly was observed to be strongest at 10 Hz stimulation frequency (Fig. S3C).

**Figure 4:**
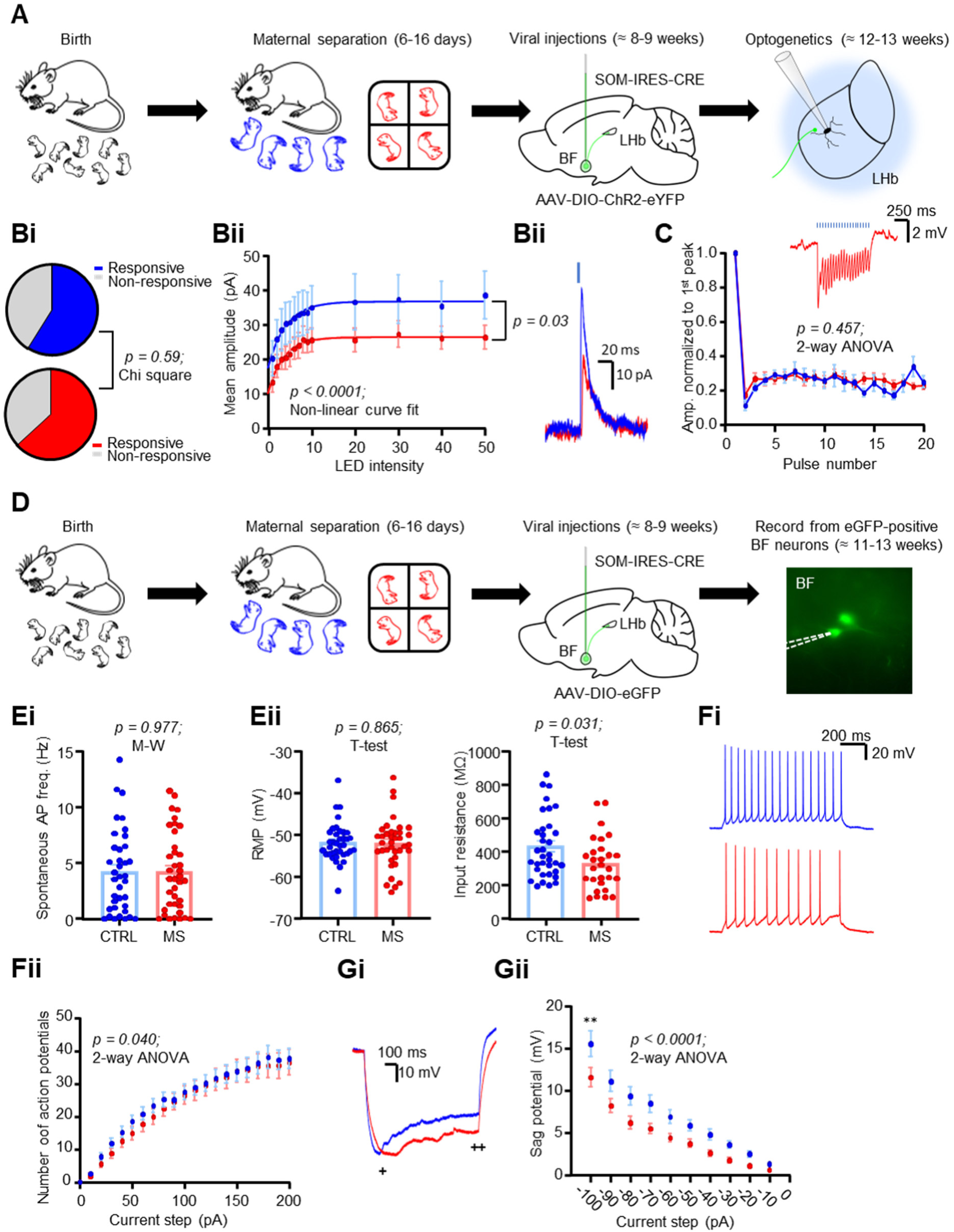
MS weakens connectivity between inhibitory SOM-positive basal forebrain neurons and the LHb. (A) Schematic illustrating experimental timeline for optogenetic experiments. (Bi) Fraction of neurons responsive to LED stimulation in both conditions. (Bii) Intensity-response curve of LED intensity plotted against mean peak amplitude of the oIPSC, with one phase exponential curve fitted for both conditions. *P* value is a comparison of plateaus from both fitted curves. (Biii) Example traces of oIPSCs from both conditions. (C) Plot of oIPSP peaks from 20 Hz LED stimulation normalised to the amplitude of the 1^st^ peak in each recording. Inset is an example recording from a neuron from an MS mouse. (D) Schematic illustrating experimental timeline for recording from putative presynaptic basal forebrain neurons. (Ei) Comparison plots of mean spontaneous activity frequency and of (Eii) passive physiological properties between neurons. (Fi) Example traces and (Fii) Input output plot of induced action potentials in response to depolarising current steps for both conditions. (Gi) Example traces and (Gii) plot of input current against sag potential in both conditions. Sag potential was calculated as the difference of the peak (+) and the steady state (++) of the membrane hyperpolarisation induced in response to hyperpolarising current steps.

Altogether, the above observations indicate a weakening of inhibitory synaptic drive from inhibitory SOM-positive pallidal forebrain neurons onto LHb neurons following MS. Due to the observed change in plateau without corresponding change in rise constant of the curve (Fig. 4Bii), and the apparent lack of change in presynaptic release probability (Fig. 4C), the case could be made that this may be due to a down-regulation of postsynaptic GABA receptors. However, this doesn’t innately rule out a change in the intrinsic excitability of the presynaptic neurons. As such, we aimed to test this by recording from presynaptic SOM-positive pallidal neurons. We first injected a retrograde Cre-dependent virus encoding tdTomato into the LHb of SOM-Cre mice. However, while we could clearly observe terminals in the LHb, and tdTomato-positive soma in the entopeduncular nucleus, where SOM-positive neurons project to the LHb, labelling in the entire basal forebrain region was very sparse (data not shown). Instead, we therefore injected a Cre-dependent anterograde virus encoding eGFP into the basal forebrain and recorded from putative presynaptic eGFP-positive neurons (Fig. 4D; n/N = 35/4 CTRL; n/N = 34/4 MS neurons / mice). These neurons were found to be spontaneously active at a similar frequency within both conditions (Fig. 4Ei; *p =* 0.977; Mann-Whitney test) and had similar resting membrane potentials (Fig. 4Eii; *p* = 0.865; unpaired T-test). However, neurons from MS mice exhibited a reduction in input resistance (Fig. 4Eii; *p* = 0.031; unpaired T-test), and a slight but statistically significant reduction in action potential firing following depolarising current injection (Fig. 4Fi and Fii; *p* = 0.040; 2-way ANOVA column factor). Additionally, these neurons exhibited a strong reduction in sag potential (Fig. 4Gi and Gii; *p* < 0.0001; 2-way ANOVA column factor). Hence, all of these observations point to a reduction in excitability in SOM-positive pallidal neurons in adult mice following MS. Altogether with our optogenetic experiments, these results indicate that MS reduces inhibitory connectivity between the pallidum and forebrain via both pre- and postsynaptic modifications.

### MS sensitises LHb neurons to acute stress

Thus far we have shown that MS induces a seemingly mild depressive phenotype within the adult mouse, which is accompanied by reduced activity within the LHb in slices and various synaptic alterations. While the changes we observed in inhibitory transmission (Figs. 3 and 4) appear to fit relatively well with the current literature, our data for the physiological properties of LHb neurons (Fig. 2) apparently goes against the central hypothesis that depression is driven by hyperactivity within the LHb. Referring again to our behavioural data, we speculated that a possible explanation for this may be that rather than being in a strongly depressed state as adults, the mice are in a state where they are only mildly depressed but rather do not respond particularly well to emotional challenge. Several observations led us to this hypothesis: firstly, the observation that the sucrose preference deficit is strongest on day 2 when the positions of the bottles are switched (Fig. 1B), which may indicate that the MS mice do not respond as well to changes in learned behaviours. Secondly, that the MS mice exhibited an increased latency to groom in the splash test (Fig. 1Dii). This is a reflection of the fact that the initial reaction of the mice to being sprayed with sucrose solution is to panic and flee, and the MS mice seemingly do this for longer and hence take longer to relax and start grooming. Thirdly, the observation that although not more spontaneously active at rest, the neurons from MS mice fire rebound bursts with a shorter latency (Fig. 2Di and Dii), possibly indicative that they are more primed to fire in response to synaptic drive.

To test this, we therefore submitted both CTRL and MS mice to an acute stressor in the form of 1 hour restraint, then immediately sacrificed them and performed acute slice recordings (Fig. 5A; n/N = 56/5 CTRL; n/N = 51/5 MS neurons / mice). As with our previous recordings, we observed no difference in passive physiological properties of LHb neurons (Fig. 5B; input resistance *p* = 0.958; T-test; RMP *p* = 0.793; Mann-Whitney test). However, here we observed a more prominent increase in intrinsic excitability in MS neurons (Fig. 5C; *p* < 0.0001; 2-way ANOVA column factor). Surprisingly, we observed a lesser difference in the latency to rebound burst, although still significant (Fig. 5D; *p* = 0.042; 2-way ANOVA column factor). As hypothesised, we found that a greater fraction of the neurons from MS mice were spontaneously active at rest following acute restraint (Fig. 5Ei and Eii; *p* < 0.0001; Chi square test). Moreover, although the average frequency of neurons which were spontaneously active was not found to be different (Fig. 5Eiii; *p* = 0.764; Mann-Whitney test), neurons from the MS mice displayed a differing distribution of spontaneous activity frequencies (Fig. 5Eiv; *p* = 0.045; Kolmogorov-Smirnov test). Interestingly, we also observed a trend towards a positive correlation between the behavioural Z score of the mice and the mean spontaneous activity frequency for each mouse (Fig. 5F; *p* = 0.063; simple linear regression), indicating that the emotional state of the mouse is a reasonably valid predictor of the response of the LHb neurons to stress. We further tested our hypothesis that the LHb neurons are more sensitive to stress histologically, by quantifying expression of the immediate early gene cFos in a separate cohort of both CTRL and MS mice either with or without exposure to acute restraint (Fig. 5Gi and Gii; n/N = 25/8 CTRL; n/N = 27/9 MS; n/N = 32/10 CTRL restraint; n/N = 31/9 MS restraint slices / mice). Restraint was able to reliably drive cFos expression in both CTRL and MS mice (Fig. 5Gi and Gii; *p* < 0.0001; one-way ANOVA), with MS mice exhibiting greater numbers of cFos-positive neurons than CTRL mice following restraint (Fig. 5Gi; *p* = 0.045; Sidak’s test) but not in non-restrained animals (Fig. 5Gi; *p* = 0.633; Sidak’s test), thus further confirming that LHb neurons in MS animals are more sensitised to stress.

**Figure 5:**
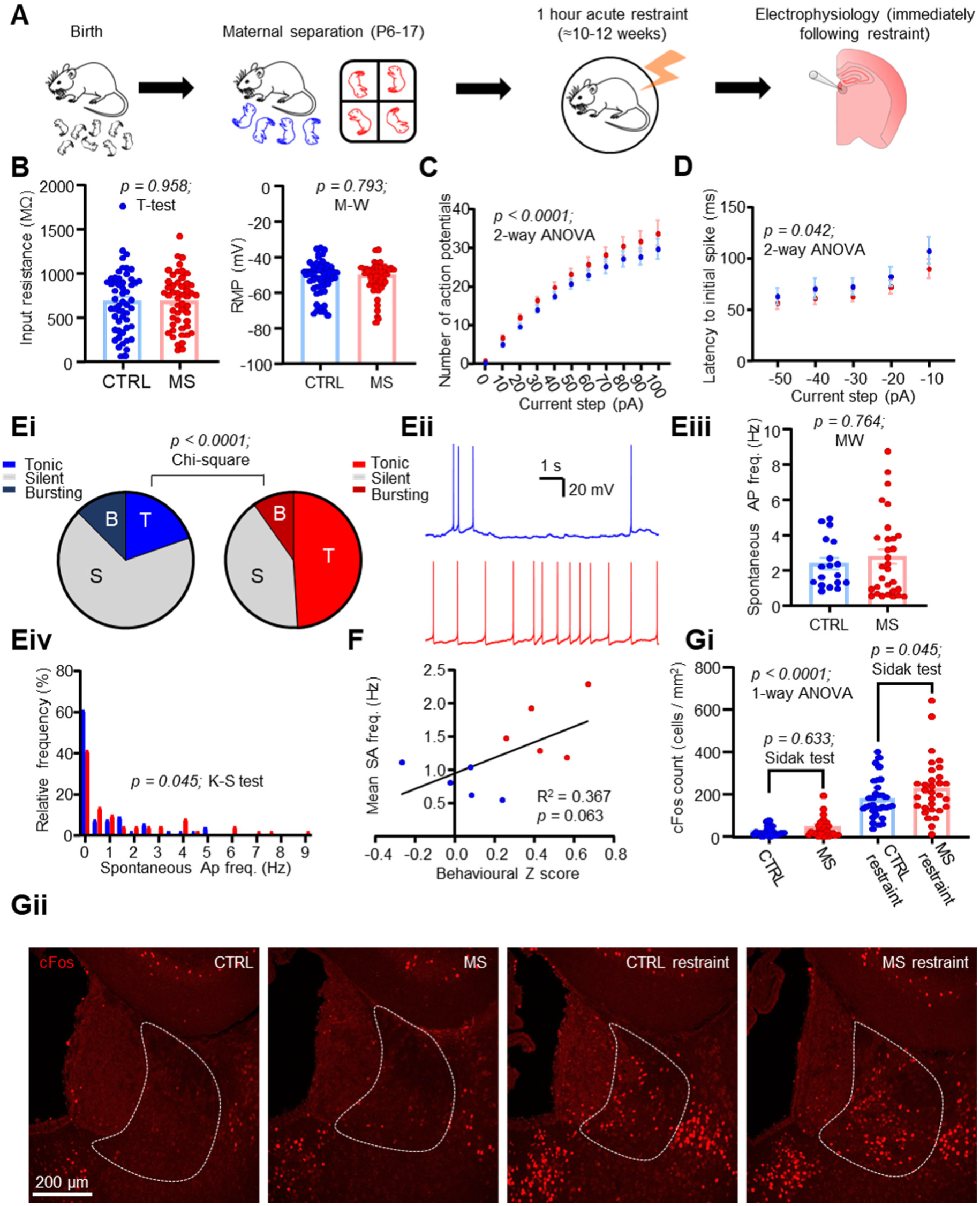
MS sensitises LHb neurons to acute restraint stress. (A) Schematic illustrating experimental timeline. (B) Comparison plots of passive physiological properties in both conditions. (C) Input-output plot of input current against mean number of induced action potentials. (D) Plot of input current against latency to initial spike of the rebound burst induced upon current step end. (E) Spontaneous activity comparison for both conditions. (Ei) Pie charts depicting fractions of recorded neurons which were classed as spontaneously active (> 0.5 Hz frequency). (Eii) Example spontaneous activity recordings from both conditions. (Eiii) Comparison plot of mean spontaneous activity frequency, for neurons which were spontaneously activity (>0.5 Hz frequency). (Eiv) Probability distribution histogram comparing mean spontaneous activity distribution for all recorded neurons between conditions. Data are 0.5 Hz bins. (F) XY plot of behavioural z score against mean spontaneous activity frequency calculated for each individual mouse recorded from. Mean spontaneous activity scores are calculated as the mean spontaneous frequency of all cells recorded from each mouse. (Gi) cFos cell counts calculated in 4 test conditions. Data are mean ± SEM of cFos counts / area calculated from individual slices. (Gii) Example confocal images of cFos immunoreactivity from each of the 4 test conditions.

Finally, we asked if this increase in activity was synaptically driven. We tested this by recording sEPSC’s in slices from CTRL and MS mice following restraint (n/N = 56/5 CTRL; n/N = 55/5 MS neurons / mice). We observed no difference in sEPSC frequency (Fig. S4A; *p* = 0.950; Mann-Whitney test) or amplitude (Fig. S4A; *p* = 0.974; Mann-Whitney test), with no obvious correlation to behavioural phenotype (Fig. S4B; *p* = 0.223; simple linear regression). However, we did observe a difference in kinetics, with many neurons in slices from CTRL mice exhibiting distinctive currents with an increased rise time (Fig. S4Ci and Cii; *p* = 0.013; Mann-Whitney test) and a strong trend towards an increased decay (Fig. S4Ci and Ciii; *p* = 0.051; Mann-Whitney test) which were less prevalent in MS mice. We also observed a clearly different data distributions for both rise time (Fig. S4Cii; *p* < 0.0001; Kolmogorov-Smirnov test) and decay (Fig. S4Ciii; *p* < 0.0001; Kolmogorov-Smirnov test). These distinctive currents appeared to be AMPA-mediated in that they were sensitive to the AMPA antagonist NBQX (Fig. S4Ci and Fig S5; *p* < 0.0001; Dunn’s multiple comparisons test), but not the NMDA antagonist AP5 (Fig. S5; *p* = 0.324; Dunn’s multiple comparisons test). A possible explanation for this may be differences in AMPA subunit composition between MS and CTRL mice, which reflects differences in ongoing plasticity (Henley and Wilkinson, 2016). Interestingly AMPA currents with extended kinetics have recently been shown to be involved in the induction of synaptic plasticity in hippocampal principal cells (Pampaloni *et al*., 2021), and as such we can speculate that this may be ongoing to a greater extent in CTRL mice, which may be related to the induction of depressive symptoms following restraint stress (Zheng *et al*., 2022). Indeed, it would make sense that these events are less prominent in MS mice, where it can be assumed that synaptic potentiation has already occurred to a greater extent. Altogether, these data are indicative of a scenario whereby LHb neurons are more responsive to stress following MS likely via differences in postsynaptic AMPA receptor subunit composition.

## Discussion

In this work we have implemented the maternal separation model of depression, in tandem with behavioural assays and *in vitro* electrophysiological recording techniques to dissect how early life stress influences the behavioural state and the underlying physiology of the LHb in adult mice. We report that the depressive phenotype is relatively weak in the adults. We also find that this is accompanied by a decrease in spontaneous neuronal activity, with a weakening of synaptic input from inhibitory SOM-positive forebrain neurons. However, rather than being more active at rest, in our model we find that the LHb neurons have a heightened sensitivity to respond to the re-exposure of stressful events. This may offer a neurobiological explanation as to why relapses occur following remission of depressive episodes.

Relating to our conclusions, arguably the most important aspect of our work that must be discussed is the validity of the MS model in successfully inducing depressive symptoms. It is important to note that historically, works implicating the MS model of depression have reported very variable results (Tractenberg *et al*., 2016). Indeed, while some studies have reported that early life stress can reliably induce depressive symptoms and drive LHb hyperactivity (Tchenio *et al*., 2017; Authement *et al*., 2018; Simmons *et al*., 2020; Langlois *et al*., 2022), others have reported a failure of early life stress to induce a depressive phenotype (Millstein and Holmes, 2007; Tan *et al*., 2017), and some evidence has even reported that it can induce resilience in the adult animals (Shi *et al*., 2021). This inherent variability in reliability of MS is likely at least partially explained by inconsistencies in protocols, with longer periods of separation generally being thought to more reliably induce aberrant phenotypes (Nylander and Roman, 2013). Another possible explanation is species and strain difference: it is believed that mice are generally more resistant to the adverse effects of MS than rats (Own and Patel, 2013; Tractenberg *et al*., 2016), and C57 mice are thought to be particularly resilient (Own and Patel, 2013). Hence, we employed the maternal separation with early weaning variant of the protocol (George *et al*., 2010), a variant of the protocol with an extended separation period (6 hours per day) and early weaning at postnatal day 17 which has previously been shown to induce a phenotype in C57 mice, and even with this optimized protocol we could only observe relatively few behavioural deficits. Importantly however, of the behavioural deficits we did observe, these all point in the direction of the MS mice exhibiting a more susceptible phenotype than the CTRL mice, hence ruling out the possibility that our model has also promoted resilience (Fig. 1).

The next key question that must be addressed is why our model induces a reduction in spontaneous activity within the LHb. It is now very well accepted that LHb hyperactivity promotes depressive behaviour (Li *et al*., 2011; Lecca *et al*., 2016; Tchenio *et al*., 2017; Cui *et al*., 2018; Yang *et al*., 2018), and our data does not superficially support this. Firstly, to address this question, we would point to the fact that in healthy animals, the LHb is active to serve an important purpose: that is to encode reward prediction error and prevent reinforcement of behaviours with negative outcomes (Hikosaka, 2010). Therefore, the relationship between LHb activity level and behavioural phenotype is likely not as simple as heightened activity equalling a more pronounced depressive phenotype, and reduced activity equalling a less depressed phenotype. Hence it may be the case that a reduction in LHb activity is also indicative of an aberrant phenotype. Indeed, a recent hypothesis has proposed that LHb hypoactivity in childhood may promote attention deficit hyperactivity disorder, which in turn primes the LHb to be more responsive to stress in adulthood (Lee and Goto, 2021). Experimental evidence has also shown that LHb inactivation induces a reversal learning deficit (Baker *et al*., 2015), which may explain why we see a prominent deficit in sucrose preference when the position of the bottles are switched (Fig. 1B). Thus, LHb hypoactivity is not likely to promote a healthy phenotype. Secondly, and in line with our first point, our recordings in CTRL mice (Fig. 2B) largely agree with the existing literature (Yang *et al*., 2018; Simmons *et al*., 2020; Langlois *et al*., 2022) in the observed distribution of tonic, bursting and silent neurons, thus supporting the claim that background LHb activity is important within healthy animals, and also ruling out the possibility of a recording artefact. Finally, although it is now well accepted that excitatory synaptic drive is potentiated onto the LHb in depression (Li *et al*., 2011), it should also be noted that there is somewhat conflicting evidence in that recent work has also observed a decrease in postsynaptic LHb AMPA receptor expression following exposure to stress (Nuno-Perez *et al*., 2021). Indeed, our data also suggest that a more depressed phenotype correlates with a reduction in spontaneous excitatory postsynaptic current frequency (Fig. S2B). Accounting for all of this evidence we would therefore conclude that in addition to LHb hyperactivity, LHb hypoactivity may also promote an aberrant phenotype and as such the reduction in spontaneous activity we observe is not mutually exclusive with the central hypothesis that LHb hyperactivity drives depression.

In terms of inhibitory signalling within the LHb, the literature is relatively consistent in that this promotes behavioural reinforcement (Faget *et al*., 2018; Stephenson-Jones *et al*., 2020; Lalive *et al*., 2022), and that inhibition of the LHb has an antidepressant effect (Winter *et al*., 2011; Huang *et al*., 2019). Consistent with this, inhibitory signalling has been shown to be perturbed in various models of depression (Shabel *et al*., 2014; Lecca *et al*., 2016; Tchenio *et al*., 2017), and indeed our data lend further support to this hypothesis. It is interesting to note that the loss of spontaneous inhibitory input we observed appears to be specific for the medial portion of the LHb (Fig. 3B). Previous work has identified a population of inhibitory LHb neurons which resides specifically within the medial LHb (Zhang *et al*., 2018; Flanigan *et al*., 2020), and hence one could speculate that activity of this population may be down-regulated following MS, therefore resulting in a loss of local inhibition. Additionally, we also report a reduction in connectivity between inhibitory pallidal neurons and the LHb, which our data also suggest involves a possible presynaptic reduction in excitability (Fig. 4E-G). While previous work has characterised inhibitory projections from various pallidal regions to the LHb relatively extensively (Golden *et al*., 2016; Faget *et al*., 2018; Stephenson-Jones *et al*., 2020; Pribiag *et al*., 2021), to our knowledge this is the first time that such a projection has been shown to be implicated in the pathogenesis of a model of depression.

## Conclusions

Depression is a complex disease, with hugely variable aetiology (Otte *et al*., 2016). Our work further complements the existing literature in that we provide evidence that LHb hypoactivity can also be associated with a depressive phenotype, and may be representative of a state where the animal is hypersensitive to stressful events. This work challenges the classical view that within the LHb, hyperactivity is the sole driver state of depressive behaviour. Further work into the specific molecular mechanisms by which these changes occur may shed new light onto the pathogenesis of depression and may unveil novel molecular targets for future therapies.

## Acknowledgement

We are grateful to the BPU staff for expert technical assistance. This work was funded by an EPSRC Doctoral Prize to JFW, a NARSAD Young Investigator Award to CW (Grant number 28217, named P&S Fund Investigator) and by the European Union’s Horizon 2020 Research and Innovation Program under Grant Agreement No. ICT-36-2020-101016787, DEEPER to CW.

## Author contributions

JFW performed the experiments, SB contributed to experiments, CW and JFW designed the study, CW supervised the work. JFW wrote manuscript with help of CW. All authors read and approved the final version of the manuscript.

## Supplemental information

**Supplementary Figure 1:**
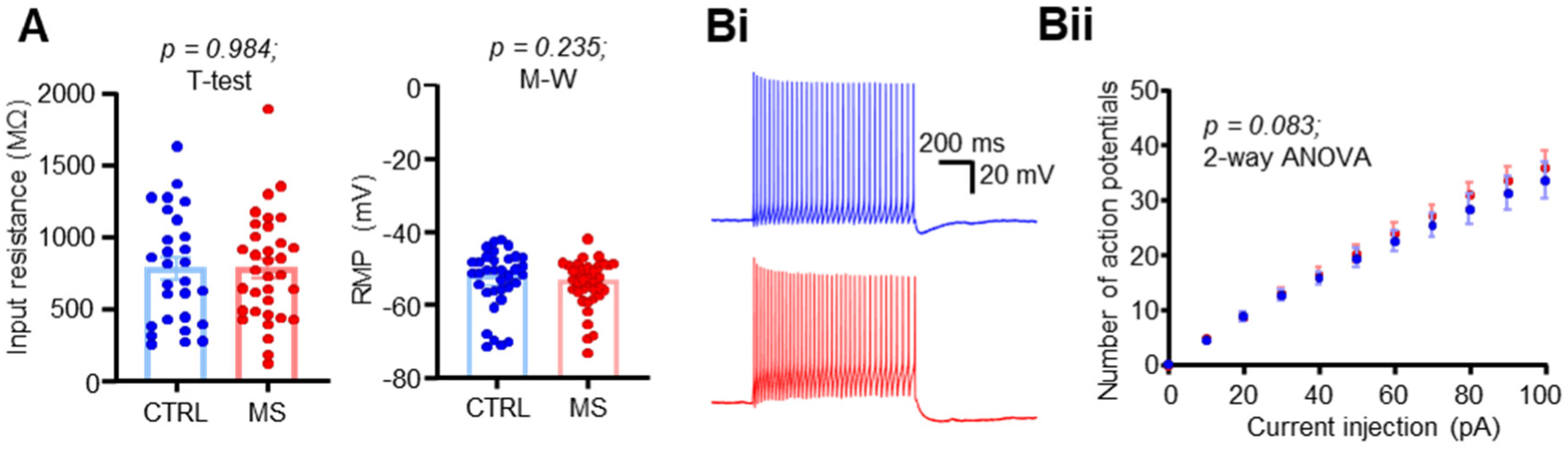
Effects of MS on passive physiological properties and intrinsic excitability. (A) Comparison plots of passive physiological properties between conditions. (Bi) Example sEPSC recordings in both conditions. (Bi) Example traces of action potential discharge in response to a 100 pA current step in both conditions. (Bii) Input-output plot of input current against mean number of induced action potentials.

**Supplementary figure 2:**
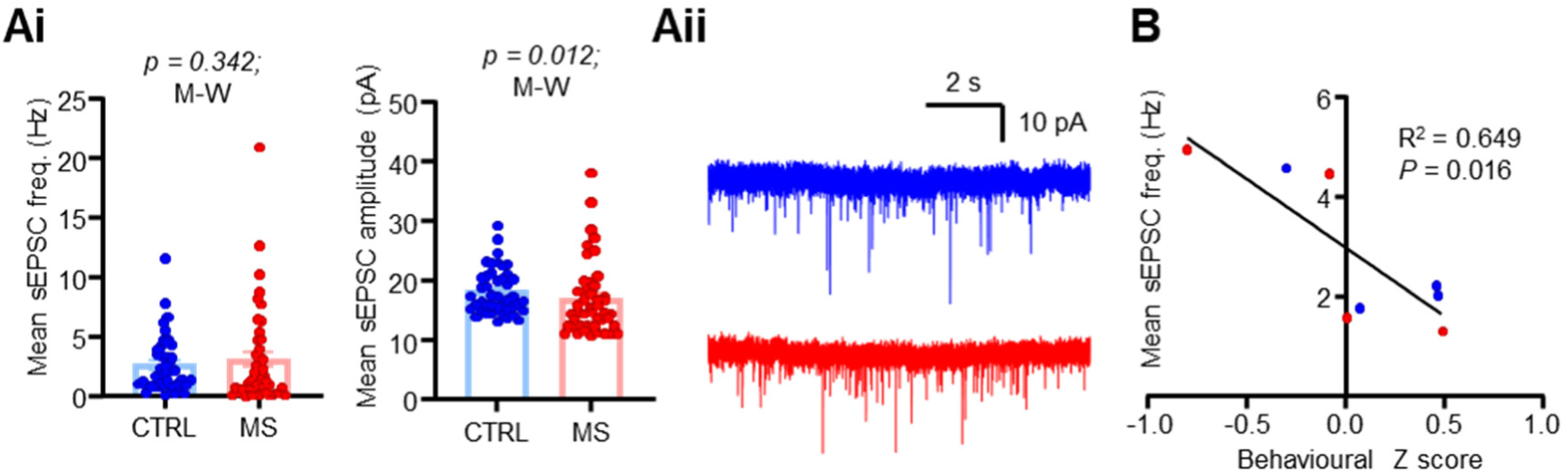
Effects of MS on spontaneous excitatory input to the LHb. (Ai) Comparison plots of sEPSC frequency and amplitude between conditions. (Aii) Example sEPSC recordings in both conditions. (B) XY plot of behavioural z score against mean sEPSC frequency calculated for each individual mouse recorded from. Mean sEPSC scores are calculated as the mean sEPSC frequency of all cells recorded from each mouse.

**Supplementary figure 3.**
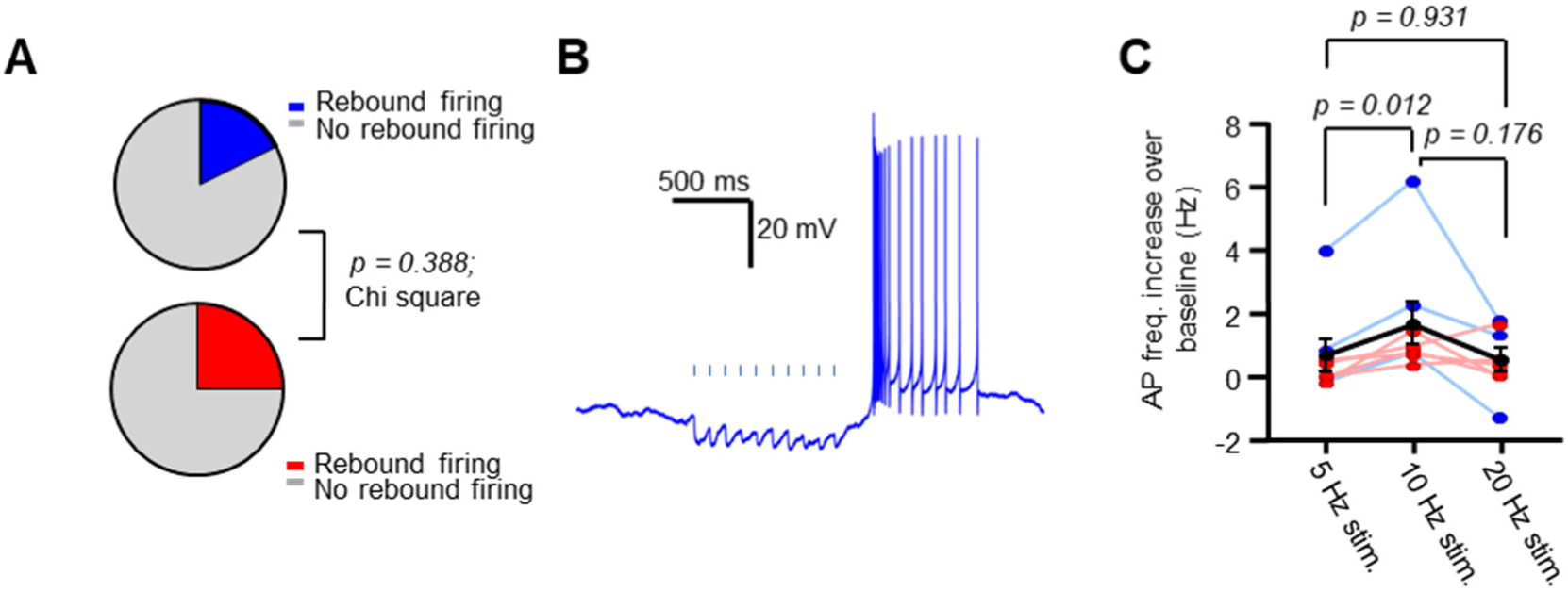
Optogenetic stimulation of inhibitory forebrain terminals can drive rebound firing within the LHb. (A) Fractions of neurons which exhibited rebound firing following optogenetic stimulation in both conditions. (B) Example trace from a neuron recorded in a CTRL mouse. (C) Comparison of rebound firing between stimulation frequencies as an increase over baseline spontaneous firing. Note here that as no difference was observed between CTRL and MS mice, these are displayed on the same plot. *p* values here are from Tukey’s multiple comparisons test.

**Supplementary figure 4:**
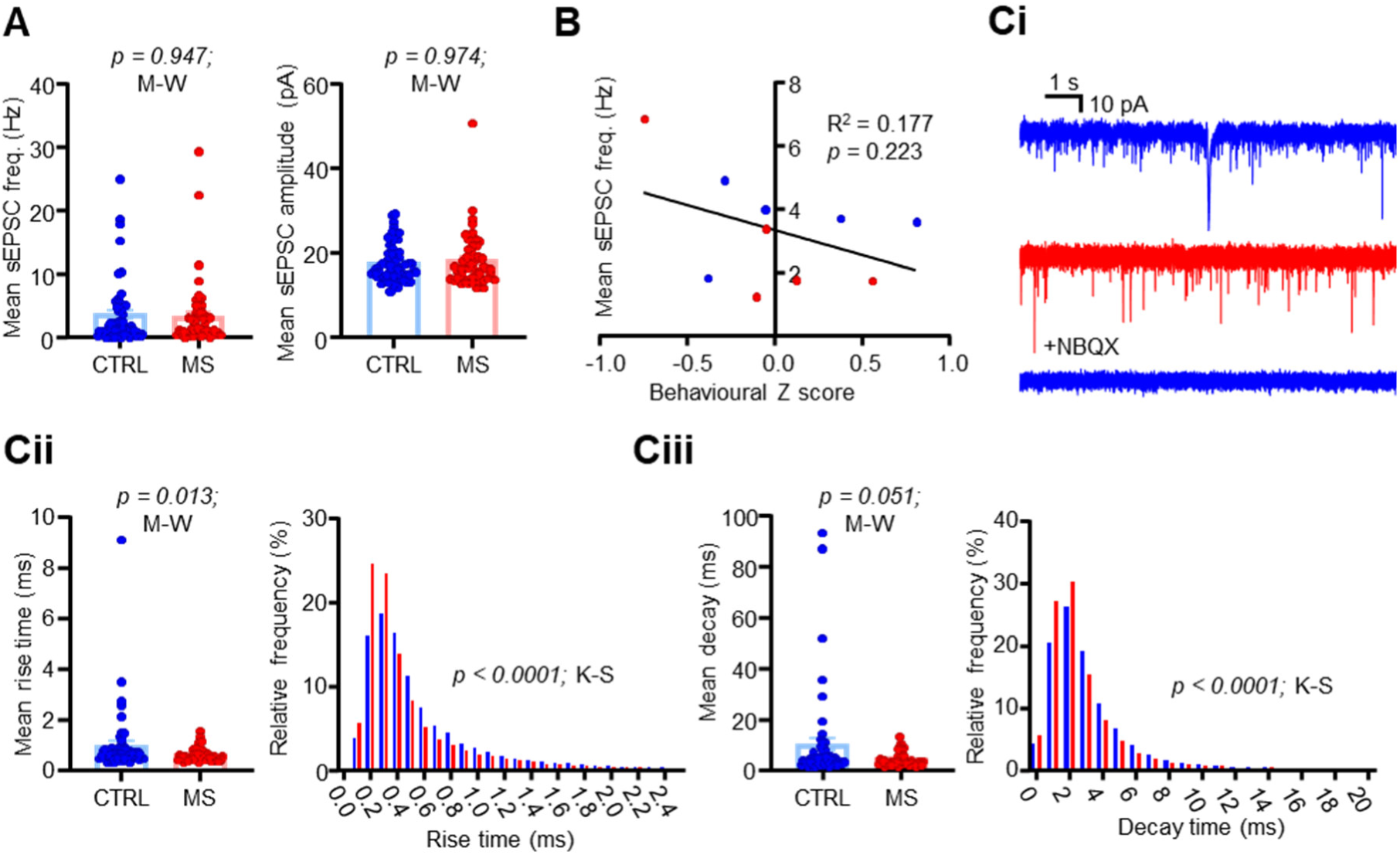
MS alters AMPA receptor signalling in response to acute stress within the LHb. (A) Comparison plots of sEPSC frequency and amplitude between conditions. (B) XY plot of behavioural z score against mean sEPSC frequency calculated for each individual mouse recorded from. Mean sEPSC scores are calculated as the mean sEPSC frequency of all cells recorded from each mouse. (Ci) Example sEPSc recordings from neurons from CTRL (top) and MS (mid) mice, and from a CTRL mouse in the presence of 10 µM NBQX (bottom). (Cii) Left: comparison plots of sEPSC rise time between conditions. Probability distribution histogram comparing mean rise time distribution for all recorded neurons between conditions. Data are 0.1 ms bins. (Ciii) As for Cii, with decay time. For probability distribution histogram, data are 1 ms bins.

**Supplementary figure 5.**
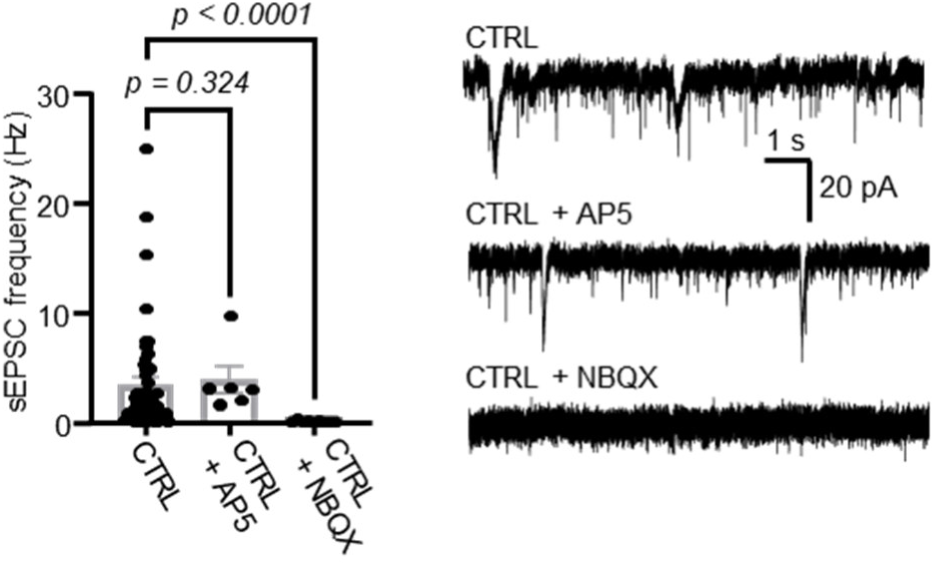
Spontaneous excitatory currents within the LHb are AMPA-mediated. Left: comparison plot of sEPSC frequencies recorded in cells without AMPA or NMDA antagonization, with 50 µM AP5 and with 10 µM NBQX. Right: example recordings from three different neurons from the same animal for each of the conditions shown in the plot on the left.

## Notes

### Competing Interest Statement

The authors have declared no competing interest.

